# Mouse Predation is Dependent on a Population of POU6F2-Positive Retinal Ganglion Cells

**DOI:** 10.64898/2026.03.16.712150

**Authors:** Fangyu Lin, Su-Ting Lin, Eldon E. Geisert

**Author notes:** Corresponding Author: Eldon E. Geisert, Professor of Ophthalmology Emory University, 1365B Clifton Road NE Atlanta GA 30322. Emails: Fangyu Lin, Su-Ting Lin, Eldon E Geisert.

## Abstract

The contribution of retinal ganglion cells (RGCs) subtypes to visually guided behavior continues to be an active area of research. We identify POU6F2-expressing RGCs essential for binocular predatory behavior. The POU6F2 RGCs are ON-OFF direction-selective RGCs that are vulnerable to glaucomatous injury. In *Pou6f2* knockout (*Pou6f2^-/-^*) mice, there is a 12% loss of RGCs, and these cells are the POU6F2-expressing RGCs. Functionally, *Pou6f2^-/-^*mice exhibited profound deficits in contrast sensitivity. In this study we found a deficit in the ability of *Pou6f2^-/-^* mice to perform a binocularly driven cricked predation test. Wildtype mice detect and capture the cricket rapidly; while, both *Pou6f2^-/-^* mice and mice with one optic nerve crushed, required significant longer times to complete the task. After optic nerve crush no further impairment in performance is seen in the knockout mouse. These data demonstrate that the POU6F2-positive RGCs are essential for this binocularly driven behavior.

## INTRODUCTION

Mice are natural predators, hunting insects and other small prey in the wild. Although laboratory mice are born and raised in controlled environment where food is provided and no hunting training occurs, they nevertheless retain this innate predatory behavior[1–3]. The predation in mice relies on multiple sensory modalities, including olfaction, audition, and vision[4–7]. Among these, vision is the primary modality driving pursuit and capture of prey[1].

Like many small ground-dwelling species, mouse eyes are laterally positioned, providing a wide panoramic field of view[5]. This broad coverage functions as a defensive adaptation, allowing mice to monitor their surroundings and detect potential predators[8]. The visual fields of both eyes overlap to form a binocular region approximately 40° wide, located directly in front of the animal[9]. Successful prey capture requires this binocular field, as it provides the depth perception and spatial accuracy necessary for rapid pursuit[10]. This behavioral reliance on binocular vision reflects the underlying connectivity of the visual pathway. Retinal ganglion cell (RGC) projections to the brain within this binocular zone are therefore critical for predatory behavior[11]. Binocular vision is essential for depth perception and precise localization of moving targets. This is demonstrated by monocular occlusion that results in a reduced ability to pursue prey and increases the time required for capture[9, 10, 12]. During prey detection, mice preferentially align targets within a stable binocular focus before initiating pursuit[5].

The mouse retina contains 46 RGC subtypes[13], and accurate hunting behavior requires coordinated activity among these diverse populations[14]. Our group has identified a novel subset of RGCs that express high levels of POU6F2. Within this population, approximately half are heavily labeled, and the remainder show moderate to light labeling[15]. We found that heavily labeled POU6F2-positive RGCs are RBPMS-positive but BRN3A-negative, and have since established that they correspond to ON-OFF directionally selective RGCs (ooDSGCs). These ooDSGCs are selectively vulnerable to damage in mouse models of glaucoma, where they are among the first RGCs to degenerate[15]. To investigate the functional role of POU6F2, we examined a *Pou6f2* knockout (*Pou6f2^-/-^*) mouse. In these mice, approximately 12% of the total RGC population are missing, corresponding to the heavily labeled POU6F2-positive RGCs[15].

Behavioral analysis revealed that *Pou6f2^-/-^*mice exhibit significant impairments in visual acuity and contrast sensitivity[15]. Based on these findings, we hypothesized that loss of POU6F2-positive RGCs would also affect other visually guided behaviors. In this study, we evaluated the ability of *Pou6f2^-/-^* mice to perform the cricket predation test[10], comparing their performance with that of wildtype littermates.

## RESULTS

### RBPMS-positive BRN3A-negative RGCs loss in *Pou6f2^-/-^* mice

To investigate the role of *Pou6f2* knockout on RGC populations, retinas were stained with two well-established RGC markers: RBPMS and BRN3A (Figure 1) and the number of labeled RGCs was quantified in *Pou6f2^+/+^* and *Pou6f2^-/-^* mice. RBPMS is a pan-RGC marker that labels the majority of RGCs regardless of subtype[16], whereas BRN3A marks a specific subset of RGCs commonly involved in image-forming pathways[17–19]. In *Pou6f2^+/+^* retinas (n=13), the average number of RBPMS-positive RGCs was 50,357, which was significantly higher (p=0.005) than the 44,322 RGCs observed in *Pou6f2^-/-^* retinas (n=4). This indicates that there is an approximate 12% loss of total RGCs in the *Pou6f2^-/-^* retina. When we quantified the number of BRN3A-positive RGCs the average number of BRN3A-positive RGCs in the *Pou6f2^+/+^* retinas (n=17) was 45,536 similar to that observed in the in *Pou6f2^-/-^* retinas 44,171 RGCs (n=7). From previous work we know that POU6F2 marks specific subgroups of RGCs, which are RBPMS-positive and BRN3A negative[20]. Thus, the 12% loss of total RGCs in the *Pou6f2^-/-^* retina is due to the specific loss of POU6F2-positive RGCs that are positive for RBPMS and negative for BRN3A.

**Figure 1.**
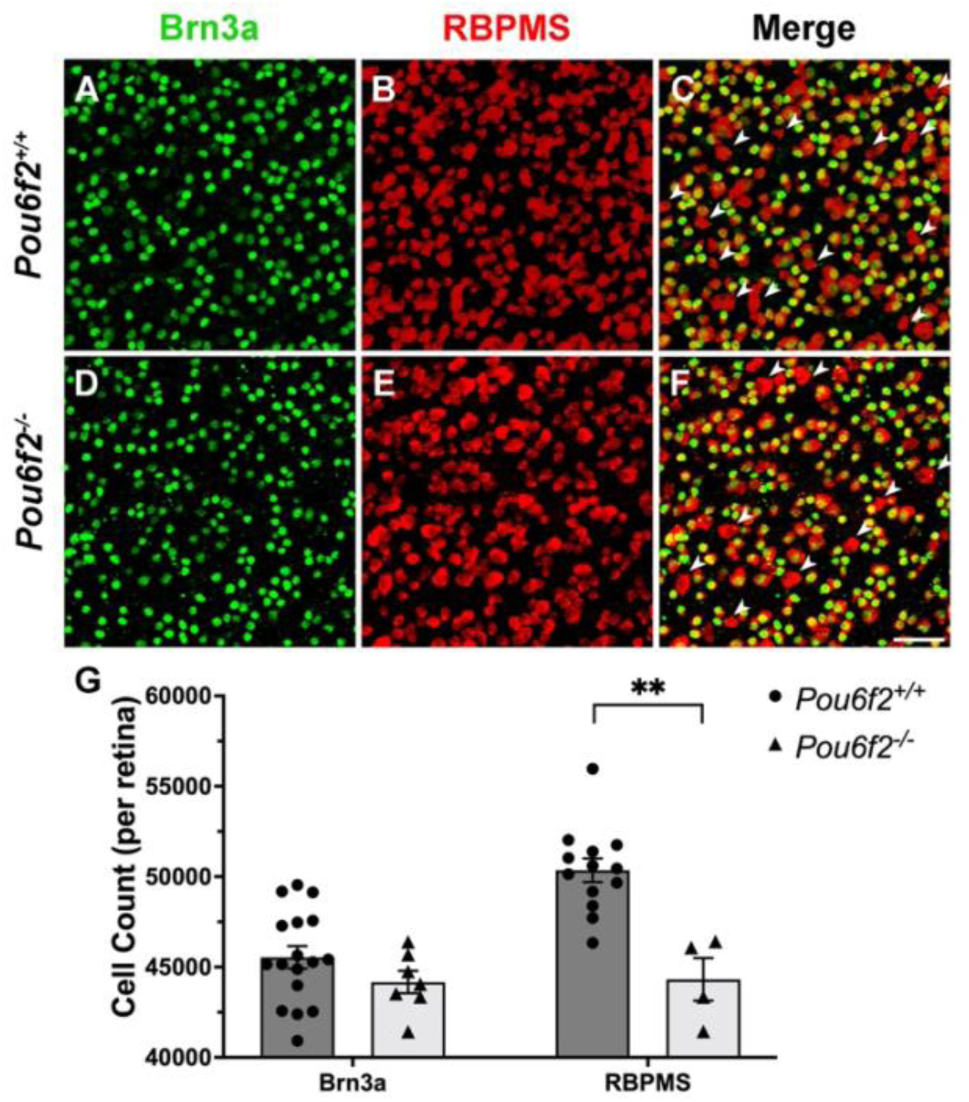
BRN3A and RBPMS Immunolabeled RGCs in *Pou6f2^+/+^* and *Pou6f2^-/-^* mouse retina. All BRN3A-positive RGCs also express RBPMS, while a subset of RBPMS-positive RGCs lack BRN3A (indicated by arrowheads in C and F). The number of labeled RGCs labeled with BRN3a is similar between the *Pou6f2^+/+^* retinas and the *Pou6f2^-/-^*retinas (A and D). There was a distinct difference in the number of RBPMS-positive RGCs within the *Pou6f2^+/+^* retinas and the *Pou6f2^-/-^*retinas (B and E) with fewer labeled cells in the *Pou6f2^-/-^* retina. When the number of RGC was quantified (G), there was no significant difference between *Pou6f2^+/+^* and *Pou6f2^-/-^*in BRN3A-positive RGCs; while there was a significantly decrease in RBPMS-positive RGCs (12%). In G, each dot or triangle represents one retina, dark grey and light grey indicate the mean RGC counts in *Pou6f2^+/+^*and *Pou6f2^-/-^* mice, respectively. These data demonstrate a loss of POU6F2-positive RGCs that are positive for RBPMS and negative the BRN3A. Scale bar = 50μm (F).

### Impaired Visual Function in *Pou6f2^-/-^* Mouse

Visually guided behavior was initially tested using OptoMotry (OMR) in both *Pou6f2*^+/+^ and *Pou6f2^-/-^* mice. Tests were performed to assess visual acuity as well as contrast sensitivity. OMR revealed no difference between *Pou6f2^+/+^*mice and C57BL/6J mice. When we examined the *Pou6f2^-/-^* mice (n=19), there was a significant depression in visual acuity relative to the *Pou6f2^+/+^*mice (n=30). On average, the *Pou6f2^-/-^* mice only reached a maximum visual acuity of 0.238 cyc/deg on average, whereas the *Pou6f2^+/+^* mice reached 0.354 cyc/deg. This difference represents an average 33% reduction in visual acuity in the *Pou6f2^-/-^* group (p<0.001, Mann-Whitney-U test, Figure 2A).

**Figure 2.**
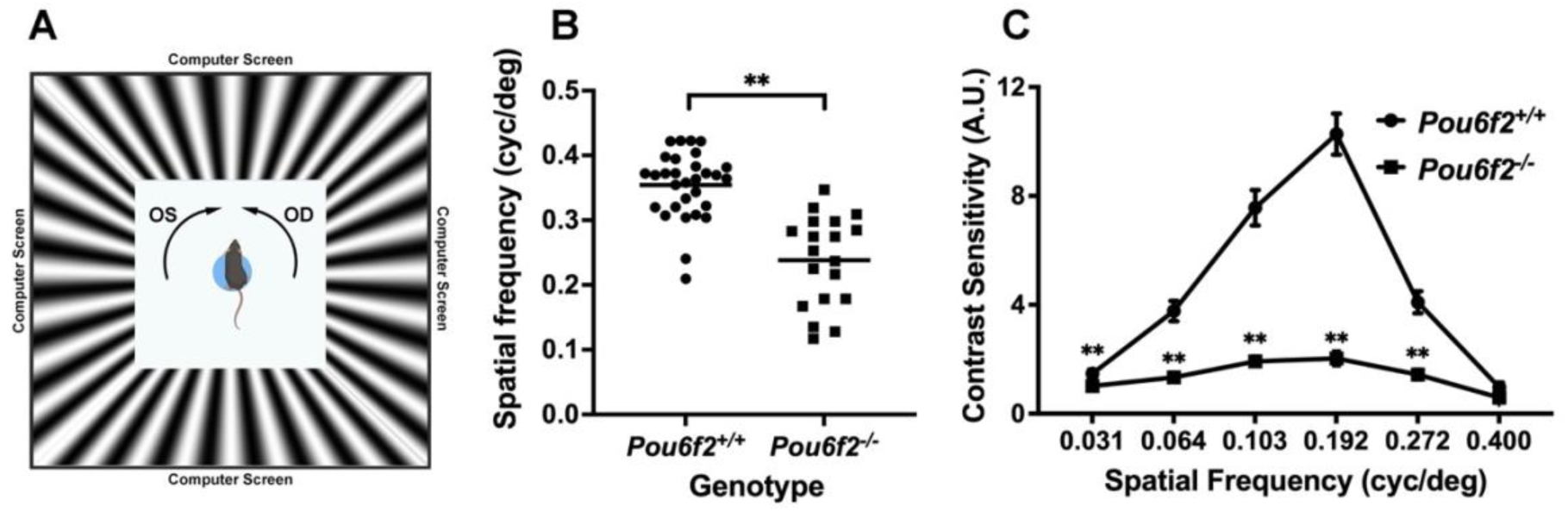
Visual acuity and contrast sensitivity assessment in *Pou6f2^+/+^* and *Pou6f2^-/-^*mice. (A) Schematic representation of the OMR system setup. The direction of rotation is randomly alternated between clockwise and counterclockwise to assess the responses of the left and right eye, respectively. The visually guided behavior was tested using the OMR to evaluate visual acuity (B) and contrast sensitivity (C) in *Pou6f2^+/+^* and *Pou6f2^-/-^* mice. (B) Individual *Pou6f2^+/+^* mice are represented by a single dot and *Pou6f2^-/-^* are represented by squares. Mice lacking *Pou6f2* showed significantly lower spatial frequency thresholds compared to wildtype mice (p<0.001). (C) Contrast sensitivity was evaluated at six spatial frequencies: 0.031, 0.064, 0.103, 0.192, 0.272 and 0.400 cycles/degree. The data is represented as means and standard error of the mean. The *Pou6f2^+/+^*mice exhibited the typical inverted U-shaped sensitivity curve, whereas *Pou6f2^-/-^* mice showed a marked reduction across all frequencies (p=0.004 at 0.031 cyc/deg, p<0.001 at 0.064, 0.103, 0.192 and 0.272 cyc/deg, p=0.519 at 0.400 cyc/deg, Mann-Whitney-U test). Both visual acuity and contrast sensitivity were significantly impaired in the knockout mice.

For contrast sensitivity, six different spatial frequencies were measured in 19 wildtype mice and 13 knockout mice (Figure 2B). Overall, the contrast sensitivity function in *Pou6f2*^+/+^ mice displayed a typical inverted U-shaped curve with a pronounced peak at 0.192 cycles/degree; while *Pou6f2*^-/-^ mouse had a markedly flattened curve, representing a significant loss of contrast sensitivity. At the lowest spatial frequencies (0.031 cyc/deg), there was small but significant difference in the wildtype mice relative to the knock-out mice (p=0.004). As spatial frequency increased, the difference became more evident. Starting at 0.064 cyc/deg, *Pou6f2*^+/+^ mice presented a sharp rise in contrast sensitivity compared to *Pou6f2*^-/-^ mice, with the greatest difference observed at 0.192 cyc/deg, where both groups reached their own peak of contrast sensitivity. At this frequency, *Pou6f2*^+/+^ mice exhibited a contrast sensitivity nearly 5-fold higher than *Pou6f2*^-/-^ mice. At high spatial frequencies (0.272 and 0.400 cyc/deg), both groups showed a decline in performance. *Pou6f2*^+/+^ mouse exhibited a steeper drop, while *Pou6f2*^-/-^ mouse maintained a relatively stable downward trend. At 0.400 cyc/deg, none of the *Pou6f2*^-/-^ mice responded, while 53% *Pou6f2*^+/+^ mice (10 out of 19) had detectable responses. This is consistent with their performance on the visual acuity task. Statistical analysis revealed significant genotype-dependent differences in contrast sensitivity at multiple spatial frequencies (p=0.004 at 0.031 cyc/deg, p<0.001 at 0.064, 0.103, 0.192 and 0.272 cyc/deg, p=0.519 at 0.400 cyc/deg, Mann-Whitney-U test). These findings demonstrate that both visual acuity and contrast sensitivity are severely impaired in *Pou6f2^-/-^* mice.

### *Pou6f2^-/-^* mice show deficits in hunting behavior

Although laboratory mouse is raised in controlled habitat throughout their lives, they retain their natural predatory instincts[3]. Mice can use both visual and auditory cues during hunting[1], but several studies have shown that they primarily employ vision to guide prey capture behavior[1, 21]. To investigate the role of visual function in hunting, we tested *Pou6f2^+/+^* (n=18) and *Pou6f2^-/-^* mice (n=14) in a prey assay across three rounds (as described in method), where live crickets were introduced into an arena and the mice were allowed to freely hunt. In this study, the group of *Pou6f2^-/-^* mice were confirmed to have visual deficit, based on previous assessments of visual acuity and contrast sensitivity. We aimed to determine whether these visual impairments would affect their hunting performance, which is an ability critical to their survival as a predatory species.

We divided the hunting sequence into three distinct phases, exploration, investigation and attempt. This framework is consistent with previous studies in laboratory mice[10, 21]. Using an overhead camera, we can detect when the mouse turned its head toward the cricket (explore), often followed by moving towards it (investigate), and finally an effort to capture it (attempt). An attempt is defined as the mouse reaching out for the cricket with its forepaws and trying to bite it. We do not consider the behavior as an attempt when the mouse was approaching and only sniffing the cricket.

The model of prey hunting in mice is predominantly vision dependent. When visual function is impaired, as seen in *Pou6f2^-/-^* mice, the predator struggles to capture its target. Data is shown in Table 2. Overall, all mice lacking *Pou6f2* gene required significantly more time (approximately 2.3 times) to complete the hunting task (average total time 40.43s for wildtype mice and 113.98s for knockout mice, respectively, p=0.014), with three knockout mice exceeding the 5-minute endpoint. From the beginning of the trial, knockout mice were noticeably slower, as demonstrated by prolonged exploration time (wildtype group 3.24s, knockout group 6.62s, p=0.044), suggesting difficulties in detecting the target. We observed that *Pou6f2^-/-^* mice often displayed signs of hesitation and reduced hunting efficiency. For example, they took longer to initiate an approach toward the cricket, and in many cases, after getting close enough to make a contact, they merely sniffed the cricket and then retreated without attempting to capture it. This contributed to an extended investigation period (wildtype group 8.82s, knockout group 80.96s, p=0.003). During the final continuous pursuit attempt, knockout mice often lost the best opportunity to successfully grab or bite the cricket. They required more time to re-engage, leading to a marked difference in the duration of the final attempt between groups (wildtype group 7.07s, knockout group 76.56s, p<0.001).

**Table 2.** Hunting behavioral differences between *Pou6f2^+/+^* and *Pou6f2^-/-^* mice before (pre) and after (post) optic nerve crush.

|  | Pre ONC |  |  | Post ONC |  |  |
| --- | --- | --- | --- | --- | --- | --- |
|  | *Pou6f2*<sup>+/+</sup> | *Pou6f2*<sup>-/-</sup> | p-value | *Pou6f2*<sup>+/+</sup> | *Pou6f2*<sup>-/-</sup> | p-value |
| n | 18 | 14 |  | 10 | 9 |  |
| Total time (s) | 40.43±10.19 | 113.98±29.20 | **0.014** | 130.48±31.05 | 155.78±36.74 | 0.744 |
| Explore (s) | 3.24±0.77 | 6.62±1.46 | **0.044** | 7.55±1.24 | 12.74±2.98 | 0.288 |
| Investigate (s) | 8.82±1.81 | 80.96±31.94 | **0.003** | 82.75±28.88 | 99.30±31.81 | 0.624 |
| Final Attempt (s) | 7.07±0.98 | 76.56±32.40 | **<0.001** | 77.96±28.96 | 109.43±39.55 | 0.568 |
| Time from First Approach (s) | 31.62±8.88 | 97.30±30.49 | **0.025** | 100.35±25.80 | 134.94±40.65 | 0.568 |

Mice were tested for cricket-hunting behavior before and after optic nerve injury. During the pre ONC phase, *Pou6f2*<sup>-/-</sup> mice spent significantly more time in all measured timepoints (including total time, exploration, investigation, final attempt, and time from first approach until successful capture) compared to *Pou6f2*<sup>+/+</sup> mice. These differences were no longer significant after ONC.

### Monocular vision reduces hunting success in the *Pou6f2^+/+^* mouse

Mouse rely heavily on vision to perceive and navigate the three-dimensional (3-D) world around them[22]. During the hunting phase, Johnson et al.[10] demonstrated that prey typically entered the binocular segment of the mouse’s visual field in front of the head, where the mouse has a best estimation of the distance and depth of the prey, giving them the best chance for a successful capture. When mice are forced to use monocular vision, they may lose the ability to accurately localize the target, resulting in a long time to complete the task.

To further explore the importance of binocular vision for hunting, we took mice from last two cohorts (total 10 *Pou6f2^+/+^* mice and 9 *Pou6f2^-/-^* mice) and performed ONC on the left optic nerve of the mice after the first cricket hunting test. In the wildtype group, ONC led to significantly increased time across all phases of the hunting task (all p<0.05), with total time, exploration time, and the interval from first approach to successful capture nearly doubling. The most dramatic changes were observed in the investigation (7-fold increase) and final attempt phases (8-fold increase). Knockout mice also showed a modest increased time in all behavior phases post-ONC, however, these increases did not reach a significant level except for a modest increase in exploration time (Table 3). The tracking route (Figure G-J) provides a clear and intuitive visualization of a representative hunting behavior. Before ONC, wildtype mice displayed more direct hunting patterns, taking the shortest path to the target. In contrast, knockout mice spent more time following and tracking the cricket, exhibiting less efficient paths. After ONC, the wildtype mice showed increased difficulties in targeting the cricket compared to their pre-ONC performance. Interestingly, we found that the post-ONC behavioral metrics in the wildtype group were close to the pre-ONC performance of knockout mice.

## DISCUSSION

In the present study, we examined the effects of knocking out *Pou6f2* on the predatory behavior of mice. Our testing was based on previous works that demonstrated the predatory behavior of mice in catching crickets in an open field[1, 10, 23]. The predatory behavior is primary driven by visual cues allowing the mice to perceive and navigate the open field to find the prey[1]. The ability of mice to rapidly detect and capture mice is dependent on binocular vison[10]. When one eye was enucleated, mice showed a profound increase in the amount of time required to capture crickets in the open field[10]. The binocular fields in mice is approximately the central 40° that is seen by both eyes, and many of the RGCs in this region are tuned to disparity[22]. Binocular vision allows for stereopsis, which is the extraction of depth information from the slightly different images received by each eye[24], as illustrated in the pole-descent cliff test[22]. This binocular input is necessary for mice to identify and approach crickets prior to capture[25]. Furthermore, Johnson et al.[10] demonstrates that binocular vision in mammals depends on ipsilateral projections (ipsi-RGCs), and selective ablation of these specific cells (<2% of RGCs) in adult mice significantly impairs their hunting success.

Three lines of reasoning from our previous study[15], led us to use this mouse predation test to examine the visually guided behavior of the *Pou6f2^-/-^* mouse relative to wildtype mice: 1) a lack of specific POU6F2-positive RGC subtypes in the *Pou6f2^-/-^*retina; 2) the fact that most of these POU6F2-positive cells are bistratified ON-OFF directionally selective RGCs; and 3) a profound deficit in visual function of the *Pou6f2^-/-^* mouse relative to wildtype mice as tested using moving gratings. When we examined the retinas of the *Pou6f2^-/-^*mouse we observed a 12% reduction in RBPMS-positive RGCs and no change in the number BRN3A-positive RGCs. These RBPMS-positive, BRN3A-negative RGCs are the heavily labeled POU6F2-positive RGCs[20]. This population of POU6F2-positive RGCs are missing in the *Pou6f2^-/-^* retina and represent several distinct subtypes of highly positive for POU6F2[15]. In the present study we stained the retinas for both RBPMS and BRN3A, replicating the loss of RBPMS-positive RGCS without any significant changes in BRN3A-positive RGCS, confirming the specific loss of POU6F2 positive RGC. Previously we demonstrated that these POU6F2-positive RGCs were bistratified and appeared to be ON-OFF directionally selective RGCs. One subset of these cells is also positive for Hoxd10 and others have demonstrated these cells are indeed ON-OFF directionally selective RGCS[15].

The *Pou6f2^-/-^* mouse also has a significant deficit in visual acuity and contrast sensitivity as defined by OMR. The OMR test is a well-established method for assessing various parameters of the visual system, particularly in the context of impaired visual function[26, 27]. It is convenient and highly reproducible. This assay is based on involuntary compensatory mechanisms of head movement triggered by motion in the surrounding environment, which serve for image stabilization on the retina during movement and enable high resolution vision[27–29]. Visual acuity provides a global measure of visual processing by both retina and visual cortex, and contrast sensitivity specifically highlights the deficits in retinal function[30]. In our study, OMR measurements in *Pou6f2^-/-^* mice revealed a significant decline in visual performance, affecting both visual acuity and contrast sensitivity.

The deficits exhibited by *Pou6f2^-/-^* mice in the OMR test prompted us to ask whether these animals also showed a reduced ability to perform visually guided tasks. One visually guided behavior is predation[11]. Hunting is a complex, visually guided behavior in mice that involves detecting the cricket, where the mouse shifts its head to bring the prey into view with both eyes, and then proceeds to directly interact with and capture the prey[5, 23]. In *Pou6f2* knockout mice, this hunting strategy was disrupted, showing increased exploration time, investigation time, final attempt time and total hunting time as compared to the wildtype mice. Throughout the process, the mice often hesitated and easily lost track of the cricket. This led to a reduced hunting success. Previously it was shown that removal of one eye also had a significant effect on the predation behavior[10]. To provide further evidence for the delayed ability to capture crickets, we performed optic nerve crush on one eye in both *Pou6f2^-/-^* and *Pou6f2^+/+^* mice. We observed that there was a significant increase in time necessary for the *Pou6f2^+/+^*mice to capture the cricket, with a 7-fold increase in investigation time and 8-fold increase in final attempt phases. By contrast, *Pou6f2^-/-^* mice showed minimal changes in the ability of performing the predation test following enucleation, and their performance was similar to the *Pou6f2^+/+^* mice that had optic nerve crushed in one eye. The impaired performance of the *Pou6f2^-/-^* mice reveals the importance of the POU6F2-positive RGCS in predatory behavior. Both Speed et al.[31] and Michaiel et al.[23] provided data that binocular clues facilitated cricket predation task.

In this study, we demonstrated that *Pou6f2^-/-^* mice exhibit impaired prey-hunting behavior under both binocular and monocular conditions, reduced optomotor responses, and significant loss of RBPMS-positive and BRN3A-negative RGCs. Together, these results suggest that the cells within the ipsilateral projecting RGCs (responsible for binocular vision and predation behavior) are the POU6F2-positive RGCs. Using the Sert-Cree mouse to eliminate ipsilaterally projecting RGCS, Johnson et al.[10] demonstrated that at least 8 different RGC subtypes contribute to the ipsilateral projection. However, since the Sert-Cree mouse does not label all ipsilateral projecting RGCs[32], other RGC subtypes might also be involved. The data from the present study indicates that POU6F2-positive RGCs are necessary for the predation behavior. In a single cell RNA seq RGC dataset[33], several RGC subtypes are *Pou6f2* positive and *Slc6a4* (Sert) positive that may be responsible for this behavior.

While we were conducting these experiments, a paper[34] from the Kerchensteiner laboratory was published that found that predation occurred without direction selectivity. The authors found that deletion of the starburst amacrine cells (which are necessary for directional selectivity of the CART-positive RGCs) resulted in a depressed response on OMR. The starburst amacrine cells were removed by a diphtheria toxin injection into SAC-DTR mice[35]. For the first days following the injection, as the amacrine cells are dying, the mice still have a normal OMR response. Once the amacrine cells die, at 10 days, the OMR response is abolished. At this time point, Krizan et al. [34] did not observe a deficit in predation behavior. These findings are in stark contrast with the observations of the present study, where in the germ-line mutation knocking out of Pou6f2 there is a deficit in the OMR response and a dramatic decline in predation behavior. At first blush, it does not appear that it is possible that both sets of data can be valid; however, that may not be the case. The first major difference is that the Krizan et al. [34] study focuses on the CART-positive RGCs that are known to receive input from starburst amacrine cells. This is a different population of ON-OFF direction-selective RGCs from the POU6F2-positive RGCs, which are CART negative. The CART-positive cells are known to have substantial projections to the lateral geniculate nucleus and superior colliculus, as well as accessary optic systems. For the POU6F2-positive RGCs, we only know about the projections of one of the subclasses of POU6F2 cells, the Hoxd10-positive cells, and they have a sparse projection to the lateral geniculate nucleus and no projection to the superior colliculus. The POU6F2-Hoxd10 RGCs do have a strong projection to the accessory optic system, including the nucleus of the optic tract. This raises the possibility that the different populations of ON-OFF direction-selective RGCs along with the specific targets in the brain may contribute to the divergent findings from the Krizan et al. [34] study and the present one.

The second major difference is that Krizan et al. [34], used a deletion in an adult mouse, while we used a germ-line mutation. Both procedures have potential artifacts associated with the method of eliminating the specific retinal cells. In germ-line mutations, there are many known complications, including developmental compensation, circuit-level reorganization, cell survival vs developmental specification, and off-target/systemic effects. Nonetheless, we observe a specific loss of RGCs in the Pou6f2-/- retina, while the remainder of the retina looks relatively normal, and we did not see any difference in ERGs between the normal mouse and the Pou6f2-/- mouse [15]. The major concern with germ-line mutations is compensation; however, in the present study there was no compensation for the predation behavior was lost. There are also concerns associated with deletion of specific cell types in a mature neuronal circuit. Deletion of the starburst amacrine cells may have unexpected consequences, including a reorganization of local circuits, inflammatory responses, glial reactivity and compensatory synaptic activity in postsynaptic neurons. In both cases, specific cells in the neuronal network are potentially being removed leaving the retina with modified circuitry that is necessary that could alter function.

In the mouse retina, many RGC subtypes recognize motion as defined by their responses to moving stimuli in their receptive fields. These motion-sensitive RGCs include ON-OFF direction-selective ganglion cells (ooDSGCs), that project to accessory optic system[36] We have found that most of the POU6F2-positive RGC are ooDSGCs which includes the Hoxd10 subtype[15]. Mice have more than 40 subtypes of RGCs, each with unique RNA profiles and morphological features[13, 37, 38]. In the macaque monkey and human, the RGC subtypes are quite different. Most of the ganglion cells are either midget RGC (70%) projecting to the parvocellular division of the lateral geniculate nucleus or parasol RGCs (20%) projecting to the magnocellular division of the lateral geniculate nucleus[13]. We found that the parasol cells are POU6F2-positive RGCs. The parasol RGCs form the magnocellular (M) pathway in rhesus macaque, which is known to play a role in image stabilization[15]. Previously we found the heavily labeled POU6F2 RGCs are among the first lost in mouse glaucoma models. This is also the case for monkey and man, where the parasol RGCs (with large diameter axons) are more severely affected by glaucoma[39, 40].

Based on the present study, if POU6F2-positive cells are absent, then there should be a deficit in visual acuity as measured by OMR. Several studies have examined the behavioral response of rodents in glaucoma models and found a decrease in visual acuity as measured by OMR[41–44]. In a bead injection model of induced glaucoma, C57BL/6 mice were subjected to 6 weeks of elevated IOP. Not only did they lose RGCs in the experimental eye they also displayed a marked depression in their OMR response[41]. In other models using Brown Norway rats, IOP was elevated by injecting NaCl solution into the episcleral veins, resulting in both increased IOP and a loss of RGCs. Visual acuity, as measured by OMR, was significantly reduced in the eyes with elevated IOP compared with the contralateral eye[43, 44]. In all these cases, investigators found that there was a decrease in visual acuity following elevated IOP. Data from the present study indicate that these deficits are potentially due to a loss of specific RGC subtypes, namely the POU6F2-positive RGCs. It is possible that a behavioral test detecting decrease in visual acuity could serve as a sensitive method for identifying the early stages of glaucoma in mice, and potentially in humans.

Overall, loss of POU6F2-positive RGCs in *Pou6f2^-/-^* mice leads to deficits in prey capture, visual acuity, and contrast sensitivity, which suggests POU6F2-positive RGCs have a meaningful impact on binocular vision and visually guided predatory behavior in mice. These results underscore the role of specific RGC subtypes in complex behaviors.

## CONCLUSIONS

*Pou6f2^-/-^* mice exhibited impaired visual function, including deficits in visual acuity and contrast sensitivity. These impairments are not attributed to BRN3A-positive RGCs, but rather to the loss of a specific subset of POU6F2-positive RGCs. The absence of *Pou6f2* may impair the ability to focus on and track moving targets, which could severely impact the animal’s effectiveness as a predator and its chances of survival in natural environments. These findings emphasize the importance of *Pou6f2* in both anatomical and functional aspects of vision.

## ACKNOWLEDGEMENT

We would like to thank Micah Chrenek for breeding and genotyping all the *Pou6f2* wildtype mice. Thanks to Prof. Fei Su and Xiaofeng Wang from Shandong First Medical University for generating tracking route from mouse prey-hunting videos. The Graphical Abstract was generated using BioRender. This study was supported by Owens Family Glaucoma Research Fund (EEG), Research to Prevent Blindness Challenge Grant, National Eye Institute Grants R01EY031042 (EEG PI) and P30EY006360 (Emory Vision Core).

## AUTHOR CONTRIBUTIONS

E.E.G. acquired funding and supervised the project. S.L. and F.L. performed the experiments. F.L. analyzed the data and wrote the first draft of the manuscript. E.E.G. reviewed and edited the manuscript. All authors read and approved the final manuscript.

## STAR METHODS

## KEY RESOURCES TABLE

| REAGENT or RESOURCE | SOURCE | IDENTIFIER |
| --- | --- | --- |
| **Antibodies** |  |  |
| Guinea pig anti-RBPMS | Millipore Sigma, Burlington | ARN1376 |
| Rabbit anti-BRN3A | Syanptic Systems | Cat.#411003 |
| Donkey anti-rabbit, Alexa Fluor 488 conjugated | Jackson ImmunoResearch | Cat. #715-545-152 |
| Donkey anti-guinea pig, Alexa Fluor 594 conjugated | Jackson ImmunoResearch | Cat. #706-585-148 |
| **Chemicals, peptides, and recombinant proteins** |  |  |
| Triton X-100 | Sigma-Aldrich | Cat. #T9284-500ML |
| Tween-20 | Fisher Bioreagents | Cat. #BP337-100 |
| Donkey Serum | Sigma-Aldrich | Cat. #S30-100ML |
| Bovine Serum Albumin | Sigma® Life Science | Cat. #A9647-100G |
| Phosphate Buffered Saline 10X Molecular Biology Grade | Corning® | Cat. #46-013-CM |
| Ketamine | Vedco - KetaVed | Cat. #VINV-KETA-0VED |
| Xylazine | Akorn - AnaSed | Cat. # 59399011150 |
| Antisedan | Zoetis | Cat. # 10000449 |
| Paraformaldehyde 32% Solution | Electron Microscopy Sciences | Cat. #15714-S |
| Proparacaine Hydrochloride Ophthalmic Solution, USP 0.5% (Sterile) | Bausch & Lomb | Cat. #24208-730-06 |
| Fluoromount-G® | Southern Biotech | Cat. #0100-01 |
| Clear Nail Polish | Electron Microscopy Sciences | Cat. #72180 |
| **Experimental Models: Organisms/Strains** |  |  |
| Mouse: *C57BL/6J* | Jackson Laboratory | Strain #: 000664 |
| Mouse: *Pou6f2<sup>-/-</sup>* | Jackson Laboratory | Strain #: 009042 |
| **Software and Algorithms** |  |  |
| Image J | NIH, USA | N/A |
| Photoshop | Adobe, USA | N/A |
| Prism 9 | https://www.graphpad.com/ | N/A |
| SPSS 24.0 | IBM | N/A |
| RGCode | https://gitlab.com/NCDRlab/rgcode | N/A |
| **Other** |  |  |
| 24-well plates | Corning | Cat. #3524 |
| 96-well plates | VWR INTERNATIONAL INC | Cat. # 41-12-21-02 |
| VWR micro cover glass | VWR INTERNATIONAL INC | Cat. #48366-227 |
| VWR micro slides | VWR INTERNATIONAL INC | Cat. #48311-703 |
| Micro dissecting forceps-Straight | Roboz Surgical Store | Cat. #RS-5070 |
| Micro dissecting forceps-Angled | Roboz Surgical Store | Cat. #RS-5095 |
| Vannas Spring Scissors-Straight | FINE SCIENCE TOOLS | Cat. #150000-08 |
| Optomotor Response | Cerebral Mechanics Inc | Model: OptoMotry |

## CONTACT FOR REAGENT AND RESOURCE SHARING

Further information and requests for resources and reagents should be directed to and will be fulfilled by the Lead Contact, Eldon Geisert.

## EXPERIMENTAL MODEL AND SUBJECT DETAILS

The *Pou6f2^-/-^* mice (B6.129-*Pou6f2tm1Nat*/J, Strain #: 009042) (knockout) and C57BL/6J mice (Strain #: 000664) were purchased from The Jackson Laboratory (Bar Harbor, ME, USA). The mice were maintained in our breeding colony. The *Pou6f2^-/-^* mice and age-matched *Pou6f2^+/+^*mice (wildtype) were produced from heterozygotic parents. All mice were 60-70 days of age at the time of experiment. The mice were housed in a pathogen-free facility at Emory University under a 12-hour light/dark cycle. Food and water were made available ad libitum until the cricket predation experimental sessions began. All animal procedures were approved by the Emory University Institutional Animal Care and Use Committee (IACUC, Protocol Number PROTO201900198) and were in accordance with the ARVO Statement for the Use of Animals in Ophthalmic and Vision Research.

## METHOD DETAILS

### Optomotor Response (OMR)

The optomotor response (OMR) system (Cerebral Mechanics Inc., Lethbridge, Alberta, Canada) uses real-time video tracking and measurement of visually evoked compensatory head movements were performed. The arena was composed of four identical, computer-controlled, interconnected 17-inch LCD screens that served as display monitors and surrounded the testing area. A 5cm-circular platform was set for the mouse to stand and move freely during testing. During OMR measurements, stimuli consisting of black and white striped patterns with differing luminosities, contrasts, spatial frequencies, and temporal frequencies were presented in a sequential or step-wise manner[26]. The optomotor stimulus rotated at a constant speed (12°/s)[28] in either a clockwise or counterclockwise direction. The direction was presented either randomly or alternated. An observer tracked the middle of the mouse head (between two ears) manually to dynamically readjust the actual physical distance to the cylinder, which guarantees that the animal perceives a constant grating when the it is moving[45]. The ability of mouse to detect and react to the stimulus by head movements is judged for each individual subject by the same observer. A minimum of three trials of head tracking observed was required to conclude that the animal could detect the stimulus. All the OMR experiments were run double-blind. Animals were coded with a number and the codes were given maintained by independent lab member (SL). The experiment was carried out by a blinded observer (FL). After the experiments were completed, the codes were released and the data was compiled.

A staircase protocol was implemented for the assessment of visual acuity (VA) and contrast sensitivity (CS) by OMR[27, 29]. VA was defined as the highest spatial frequency that the mouse could track in either direction. During VA testing, the sinusoidal gratings were all at 100% contrast. Based on the staircase procedure, spatial frequency was systematically increased or decreased depending on the mouse’s response. After 10 reversals around the highest detectable spatial frequency, the data points from these reversals were averaged and recorded as the VA. For CS testing, a fixed spatial frequency was selected, and the stimulus began with a grating of 100% contrast, which was then systematically reduced until the contrast threshold was identified. A staircase protocol with 10 reversals was also applied for CS measurements. The threshold was calculated as a Michelson contrast based on the screen’s luminance (maximum - minimum)/(maximum + minimum)[27]. The contrast sensitivity plotted on the graph was the reciprocal of the threshold. In our study, contrast thresholds were determined at six spatial frequencies (0.031, 0.064, 0.103, 0.192, 0.272, 0.400 cyc/deg).

### Cricket-predation Test

Forty-eight hours prior to the start of training, mice were individually housed, and three crickets were introduced to each cage with food pellets daily. Food pellets were removed 16-18 hours before the first training session, and three crickets were provided to each mouse. Training was conducted for 4 consecutive days in the behavioral arena (Length: 40cm, width: 30cm, height: 30cm, Figure 3A). On the first day of training, mice were placed in the arena to acclimate for 3 minutes. Then a cricket was introduced into the center of the arena. Mice were given up to 5 minutes to capture the cricket. After either successful capture or the 5-minutes limit, the arena was cleaned, and a new cricket was released. This procedure was repeated three times for each mouse. After daily training, mice were returned to their home cages and provided access to food pellet for 4-5 hours, after which the pellets were removed overnight, with three crickets remaining in each cage. Training continued for four days, followed by a test session on the fifth day. During the test, a cricket was introduced into the arena, the behavior was recorded using an overhead camera when a cricket was introduced into the arena. The experiments were run on three different cohorts of mice. The second and third group of mice were retested 3 days after ONC.

**Figure 3.**
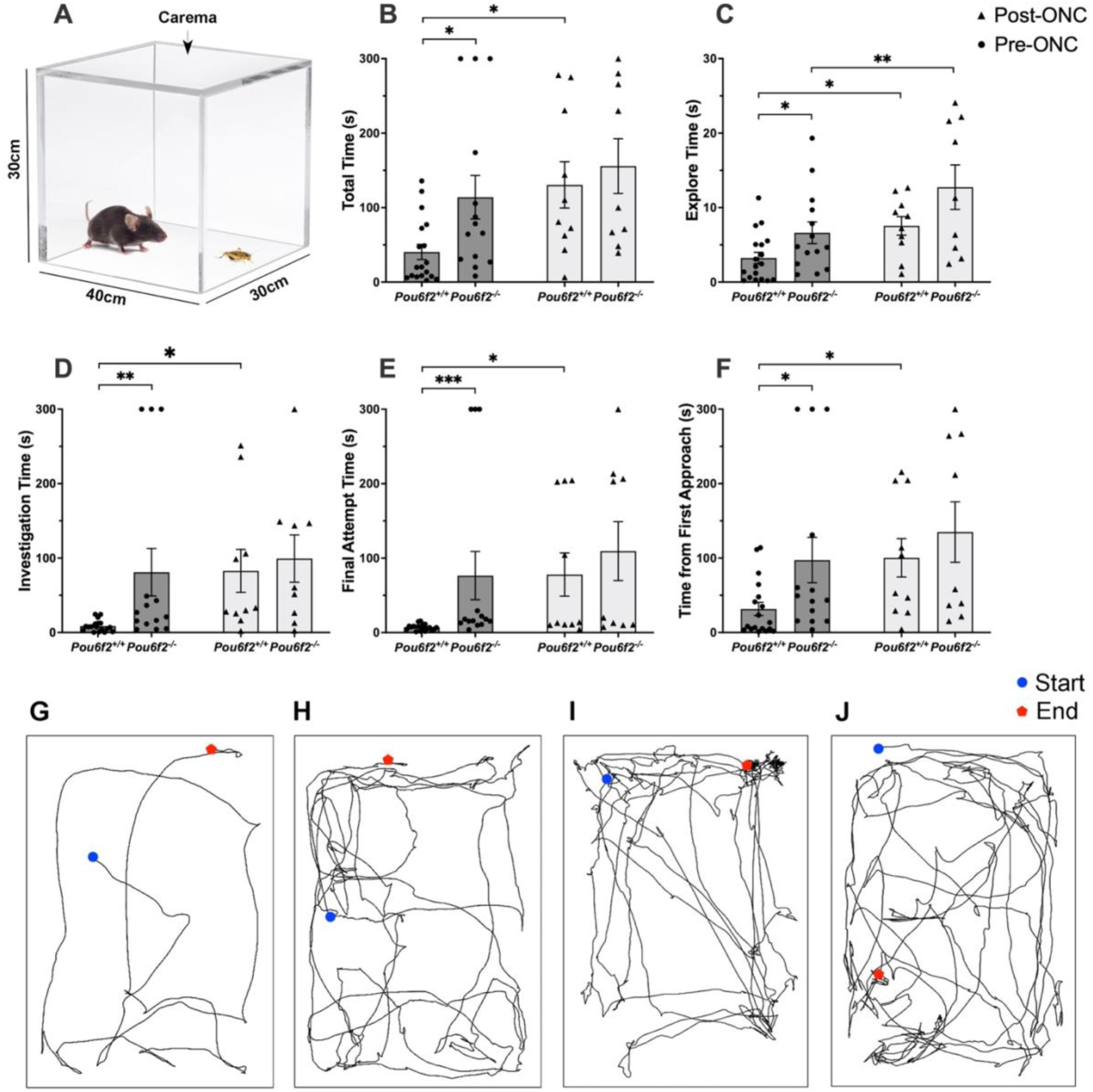
Mouse predation test including cricket capture times and tracking traces of mice before and after ONC. The arena for tracking predator-prey interactions is shown in A. B through F display quantitative differences in predation behavior (all mice in three rounds). (B) Total time: the time from the introduction of a cricket to the mouse’s successful capture. (C) Explore, time from the introduction of a cricket to the mouse’s first orientation toward the cricket. (D) Investigate, the time from the introduction of a cricket to the mouse’s first approach. (E) Final Attempt: the time from the mouse’s final approach to its successful capture. (F) Time from First Approach, the time from the mouse’s first approach to its successful capture. In all graphs, each dot represents the average of three trials for a single mouse. Data is shown for *Pou6f2^+/+^* and *Pou6f2^-/-^* mice before (Pre-ONC) and after optic nerve crush (Post-ONC). Data is shown in mean ± SEM. Significance was *p < 0.05, **p < 0.01, or ***p < 0.001 using a Mann-Whitney U test. G-J is representative overhead tracking traces of mice capturing a cricket: *Pou6f2^+/+^* mouse before ONC (G), *Pou6f^-/-^*mouse before ONC (H), *Pou6f2^+/+^* mouse after ONC (I), and *Pou6f^-/-^* mouse after ONC (J). Blue circle: the starting position of the mouse when the cricket was introduced into the arena. Red pentagon represents the end point, either end with successful capture of the cricket or the endpoint of the 5-minute trial.

**Figure 4.**
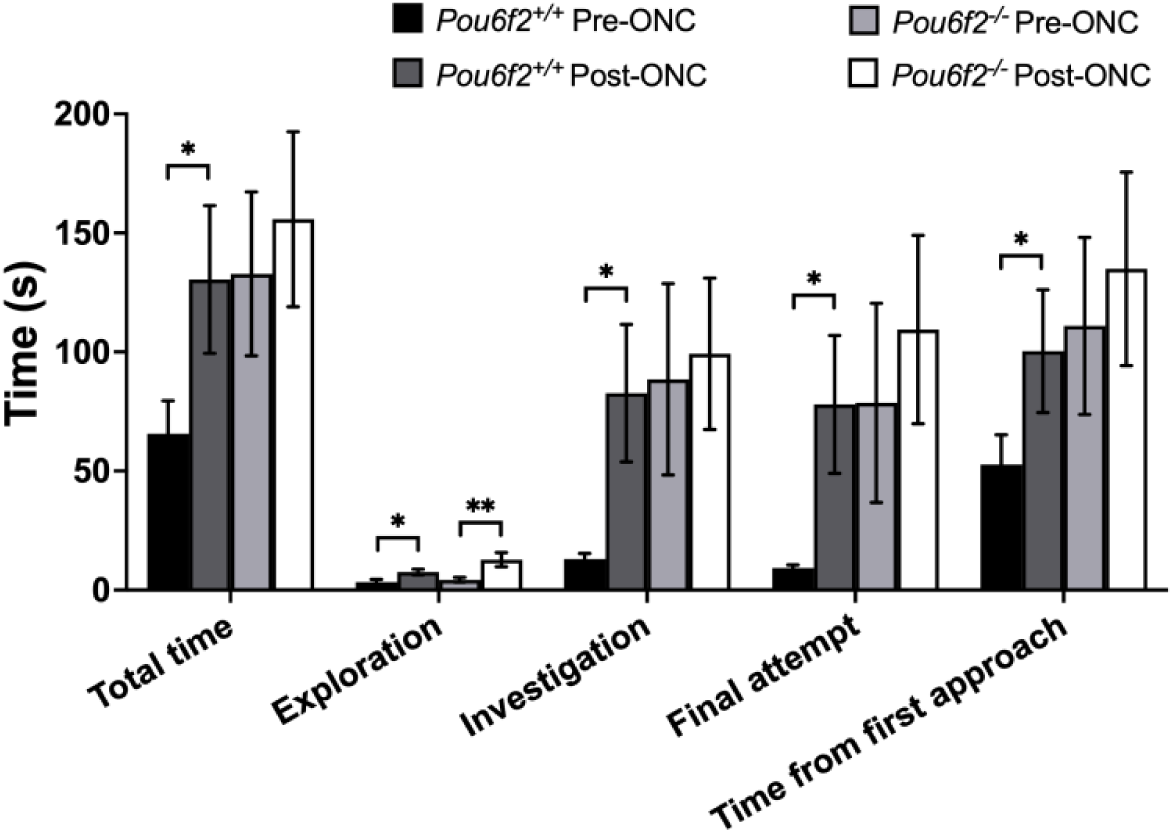
Quantitative differences in predation behavior pre- and post-ONC within each genotype in the last two rounds of mice. Following ONC, *Pou6f^+/+^* mice showed significantly increased durations across all measured timepoints (including total time, exploration, investigation, final attempt, and time from first approach until successful capture, all p<0.05), indicating impairment of visually guided behavior. In contrast, *Pou6f2^-/-^*mice displayed a significant increase only in exploration time (p=0.008), with no significant changes observed in the other measures after ONC (all p>0.05). And post-ONC behavioral metrics of *Pou6f^+/+^* mice were close to the pre-ONC performance of *Pou6f2^-/-^* mice (all p>0.05, Mann-Whitney U test). Data is shown in mean ± SEM. Significance was either *p < 0.05, **p < 0.01, using a paired-T test.

Videos were independently reviewed by two blinded researchers (FL and SL), and averages of the measurements were calculated. Prior to analysis the videos, the following behavioral timepoints were defined and agreed.

1. Orient: The moment when the mouse turns its head toward the cricket.
2. Approach: The mouse begins moving toward the cricket and makes physical contact, attempting to capture it.
3. Successful Capture: The mouse grabs the cricket with its hands or mouth and hold onto it briefly, either consuming the cricket or walking away with it.

Behavioral parameters measured in this study:

1. Total time: The duration from the introduction of the cricket into the arena to the mouse’s successful capture, or 5 minutes if no capture occurred.
2. Explore: Time from a cricket is introduced into the arena to the mouse’s first orientation toward the cricket.
3. Investigate: Time from a cricket is introduced into the arena to the mouse’s first approach.
4. Final Attempt: Time from the final approach to the moment of successful capture.
5. Time from First Approach: Time from first approach to the moment of successful capture.

### Optic Nerve Crush (ONC)

Optic nerve crush was performed as previously described by Templeton JP[46] and Jiaxing W[47]. Mice were deeply anesthetized with intraperitoneal injection of 100mg/kg ketamine (Cat. #VINV-KETA-0VED, Vedco Inc., Saint Joseph, MO) and 10 mg/kg xylazine (AnaSed; NCD 46066-750-02, Pivetal, Greeley, CO), then 0.5% proparacaine eye drops (Cat. #24208-730-06, Bausch and Lomb, Rochester, NY) was used as local anesthesia. Under the binocular operating microscope, a small incision was made approximately 2mm lateral to the conjunctiva of the left eye. A fine pair of forceps (Cat. #RS-5005, Roboz Surgical Store, Gaithersburg, MD) was used to gently grasp the edge of the conjunctiva adjacent to the globe, and the eye was pulled out carefully and rotated laterally. Then the posterior aspect of the eye was exposed and the optic nerve was able to be observed. The surrounding connective tissue and muscle were carefully separated from the optic nerve. The exposed optic nerve was then clamped with a curved crossover tweezer (No. 11487-11, Fine Surgical Instruments, Hempstead, NY) for 5 seconds. Following the procedure, the forceps were removed, and the eye was allowed to return to its natural position. No sutures were required. Triple antibiotic ointment (NDC# 24208-0780-55, Bausch & Lomb, Bridgewater, NJ) was applied to the surgical site. Mice received 0.5 mg/kg buprenorphine SR (Wedgewood Pharmacy) immediately following the surgery. Once they were fully recovered, the mice were returned to a clean cage and monitored daily for 3 days postoperatively. If excessive bleeding occurs during the surgery, the mice were immediately euthanized. Mice showing signs of pain or distress were also euthanized. All the intravitreal injections and ONC procedures were performed by a well-trained postdoctoral fellow (FL) to minimize technical variability.

### Immunohistochemistry

After completing all the experiments procedures (OMR, ONC and cricket-predation test), the mice were anesthetized as previously described, followed by transcardial perfusion with phosphate-buffered saline (PBS) and then 4% paraformaldehyde in PBS (pH 7.3). The eyes were removed, and the retinas were carefully dissected and flattened into a cloverleaf shape with four quadrants[20]. Retinas were blocked in a blocking buffer containing 3% bovine serum albumin and 3% donkey serum in 0.5% Triton X-100 in PBS. Subsequently, the retinas were incubated in primary antibodies for over two nights at 4℃. After incubation, the retinas were rinsed three times with PBST (0.1% Tween-20 in PBS) at room temperature for 15 minutes each. The retinas were then placed in secondary antibodies overnight at 4℃, followed by three rinses with PBST. Finally, the retinas were mounted on glass slides and cover slipped using Fluoromount-G (Cat. #0100-01, SouthernBiotech, Birmingham, AL) and prepared for imaging. All antibodies and their concentration used are listed in KEY RESOURCES TABLE.

### Image processing

Retinal flat mounts were imaged using a Nikon Ti2 confocal microscope equipped with an A1R confocal imaging system (Nikon, Inc., Melville, NY, United States) under 40X magnification. Z-stacked images were taken at 0.1μm intervals, with a total of 300-600 optical slices per retina to capture the entire area.

### Automated Counting

The total number of RGCs labeled with RBPMS and BRN3A across the entire retina was quantified using the deep learning-based tool RGCode, as previously described[20, 48].

### Statistical Analysis

Data are presented as mean ± standard error of the mean (SEM). Differences between genotypes before and after ONC were analyzed using the Mann-Whitney U-test. Within each genotype group, differences between before and after ONC were analyzed using paired t-tests. All statistical analyses were performed using SPSS statistics package 24.0 (SPSS, IBM, Chicago, IL, United States). A p-value of less than 0.05 was considered statistically significant.

